# Optimising multi-batch TMT analysis to mitigate inflation of missing values, false positives and diminished inter batch accuracy

**DOI:** 10.1101/497396

**Authors:** Alejandro Brenes, Jens Hukelmann, Dalila Bensaddek, Angus I. Lamond

## Abstract

Multiplexing strategies for large-scale proteomic analyses have become increasingly prevalent, TMT in particular. Here we used a large iPSC proteomic experiment with twenty-four 10-plex TMT batches to evaluate the effect of integrating multiple TMT batches within a single analysis. We reveal a significant inflation rate of missing protein and peptide values and show that precision decreases as multiple batches are integrated. Additionally, we explore the effect of false positives using Y chromosome specific peptides as an internal control to quantify the effect of co-isolation interference, as well as primary and secondary reporter ion interference. Based on the results we suggest solutions to mitigate these effects. We show using a reference line can increase precision by normalising the quantification across batches and we propose experimental designs that minimise the effect of cross population reporter ion interference.

## Main text

High-throughput, shotgun proteomics, using data dependent acquisition (DDA), now enables the comprehensive study of proteomes, allowing the identification of 10,000 or more proteins from cells and tissues^1–3^. However, to achieve such deep proteome coverage using DDA, extensive prefractionation of extracts prior to mass spectrometry (MS) analysis is frequently required^1,4^. To evaluate statistically the significance of the resulting data, a minimum of 3 replicates for each sample/condition is also necessary^5,6^. The data acquisition time involved is increased still further for experiments that analyse the multi-dimensional characteristics of the proteome; e.g. studying differences in protein subcellular localisation, turnover rates, post-translational modifications (PTMs) and protein-protein interactions^7^.

To cope with the challenges of large-scale proteomics analyses, strategies have been developed to allow multiple samples to be analysed in parallel, through multiplexing isotopically tagged proteins ^8,9^. The most widely used MS multiplexing methods, TMT^10^ and iTRAQ^11^, use isobaric tags for simultaneous peptide identification and quantification. TMT in particular has increased in popularity and is now widely used ^12,13^. This reflects the ability of multiplexed TMT to increase sample throughput in proteomics studies and reduce the “missing values” problem that arises from the stochastic sampling inherent in DDA proteomics ^14,15^. Thus, within a single multiplex TMT batch, the number of missing values at the protein level is low, frequently <2%^12^. Furthermore, the precision of the quantification within a multiplex TMT batch is high^16^. However, it is less clear how well multiplex TMT performs for very large-scale analyses, involving many multiplex-TMT batches.

In this manuscript, we analyse a recent data set from a proteomic study of human iPSC cells, involving 24 separate 10-plex TMT batches^17^. We compare the quantitation of data both within and between separate 10-plex batches and focus our analysis on 3 main issues: (i) missing values, (ii) accuracy of quantification and (iii) the effect of both reporter ion interference (RII) and co-isolation interference (CII).

We show an inflationary effect on missing values as data from multiple batches are integrated both at the protein and peptide level. We evaluated reproducibility both by studying the coefficient of variation (COV) within each 10-plex TMT batch, and by comparing technical replicates that were common to every batch. Finally, we explored the effect of co-isolation and reporter ion interference, which can both reduce quantitative accuracy. For this, we identified peptides matched uniquely to Y chromosome genes to provide an internal control, which was used to evaluate the presence of false positive peptide signals between samples derived from male and female donors.

## Results

### Missing values in TMT

A known advantage of using multiplex TMT analysis is the low index of missing values that are present within a single TMT batch. Recent studies report as low as <1% missing values at the protein level ^16^, albeit data are usually not reported at the peptide level.

We started by analysing the iPSC 10-plex TMT data for the number of missing values at the protein level within each TMT batch (Fig. 1a). The preliminary results are consistent with previous reports, i.e. 79% of the 24 different 10-plex TMT batches show <1% missing values at the protein level, with only 1 outlier with missing protein values >2%. Furthermore, when we analyse the data at the peptide level, there is very close agreement to the protein data, with 79% of the 24 different 10-plex TMT batches having <2.5% missing peptide values.

**Figure 1.**
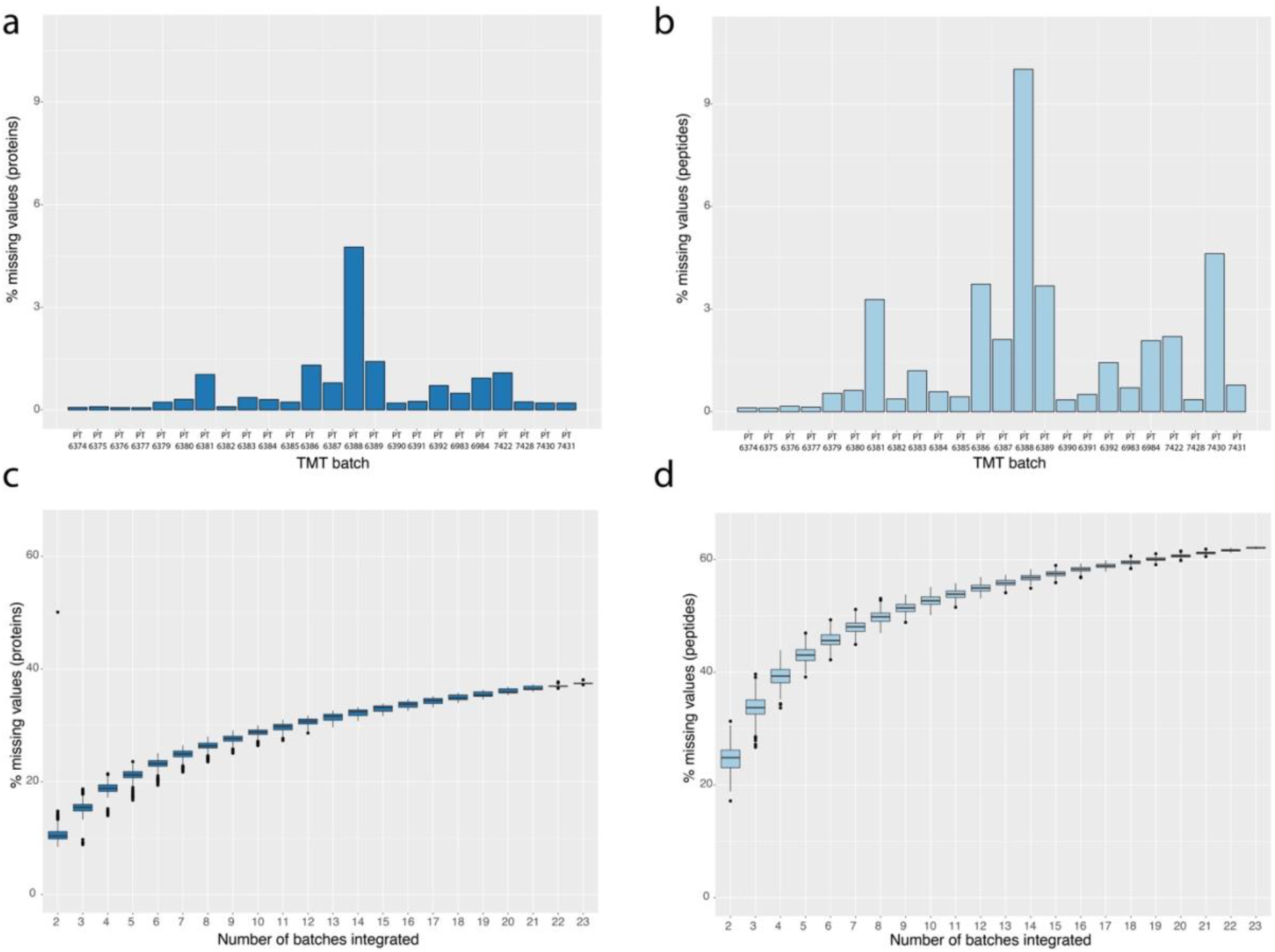
Protein and peptide missing values: (a) Percentage of missing values for each TMT batch calculated at the protein level. (b) Percentage of missing values for each TMT batch calculated at the peptide level. (c) Box plot showing the results for protein missing values as a function of the number of 10-plex TMT batches. (d) Box plot showing the results for peptide missing values as a function of the number of 10-plex TMT batches.

However, these results do not address the effect of integrating data from multiple, independent 10-plex TMT batches into a single analysis. To study the effect of data integration, we increased the number of batches selected, from 2 to 23 and recalculated the number of missing values that were present (Fig. 1c&d; see methods). At the protein level, the median number of missing values increases from 0.28% with one 10-plex TMT batch, to 10.53% when data from a second 10-plex TMT batch were integrated (Fig. 1c). When we integrate data from 12 different 10-plex TMT batches, the median number of missing values at the protein level expanded to >30%. For example, even some highly expressed histones (HIST2H2AB, 86^th^ percentile of abundance), are only detected in 66% of the cell lines.

This situation was exacerbated when the analysis was performed at the peptide level (Fig. 1d). When integrating data from just two 10-plex TMT batches, the median number of missing peptide values was >24%. Even more striking, it only required integrating data from 8 different 10-plex TMT batches to produce ~50% missing values at the peptide level.

Based upon these results, we decided to perform a more in-depth analysis on the inflation rate of peptide missing values. We observed that the number of peptides identified within each 10-plex TMT batch is relatively constant (Fig. 2a), but quite variable across different batches. Thus, we identified one outlier batch amongst the 24 batches showing TMT channels with fewer than 60,000 peptides identified, while the median number of peptides identified per batch was 93,140 with a standard deviation of 13,403. To further analyse these peptide level data, we first median-normalised the MS3 intensities for all peptides in all cell lines (see methods). The median values across the median-normalised log10 MS3 intensities spanned 6 orders of magnitude (Fig. 2b).

**Figure 2.**
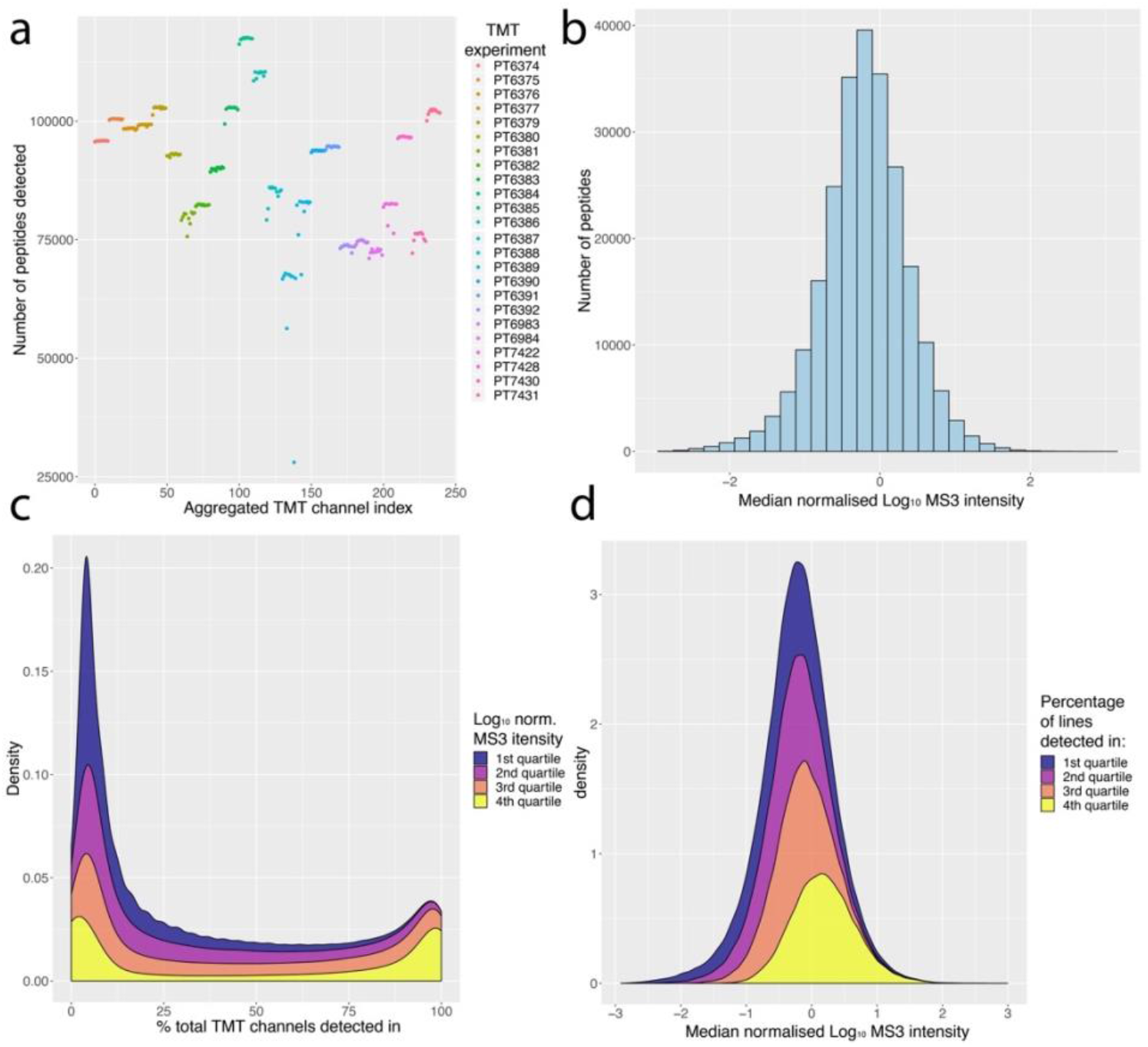
Peptide identifications and intensities: (a) Number of peptides identified per TMT channel, coloured by TMT batch. (b) Histogram of the median, median-normalised log10 MS3 intensity for each peptide. (c) Stacked density plot showing quartiles of the median log_10_ normalised MS3 intensity and percentage of cell lines they are detected in. (d) Stacked density plot showing quartiles of the percentage of cell lines in which each peptide is detected and their corresponding median log_10_ normalised MS3 intensity.

We next analysed the peptide dataset by quartiles, based on the normalised log_10_ MS3 intensity values (Fig. 2c). The first quartile represented the 25% least abundant peptides and the fourth quartile the 25% most abundant peptides. There are no peptides within the first quartile that are detected in all TMT channels and only 10 peptides that are seen in > 99% of the TMT channels. As DDA selects the n most abundant ions reaching the mass-spectrometer during a MS1 scan^18^ (where n typically is 10-30), this bias is predictable. However, when we analyse the results from the fourth quartile (25% most abundant peptides), we see that 44% of these peptides are still detected in <25% of the TMT channels and only 11% of these peptides were detected in all the TMT channels. There are 9,187 peptides that were detected in all TMT channels, and their minimum normalised intensity, −0.33, is almost equal to the overall median of the distribution, −0.20.

Next, we analysed the data by comparing the identification quartiles, organised by the percentage of TMT channels in which they were detected (Fig. 2d). The first quartile represented the 25% of peptides that were detected least frequently, i.e. in less than 11 TMT channels (see supplemental data). Of these peptides >29% had a median-normalised log^10^ MS3 intensity higher than the distribution median (−0.20), highlighting that even relatively abundant peptides are not identified consistently. Overall, ~41% of all peptides are detected in <10% of all TMT channels, while ~50% of peptides are detected in <20% of all TMT channels.

### Variation between 10-plex TMT batches

Multiple studies have documented TMT as a method producing precise quantitation, in some cases having a coefficient of variation (COV) ~3x lower than comparable label free data^16^. Most of these studies have focused on analysing quantitative precision within a single multiplex TMT batch, and do not explore the effect of integrating data from multiple TMT batches into one analysis. However, projects involving large scale proteomic analyses of multiple cell lines and/or conditions, need to employ multiple TMT batches in a single experiment^8^.

We calculated protein copy numbers for 216 different iPSCs lines, and 24 technical replicates of a single, control iPSC line, across 24 separate 10-plex TMT batches^17^. We then proceeded to calculate Lin’s concordance correlation coefficient^19^ for every iPSC line within each TMT 10-plex batch, and for all the technical replicates of the control line, channel TMT^10^ 126, across all the 24 10-plex TMT batches (Fig. 3).

**Figure 3.**
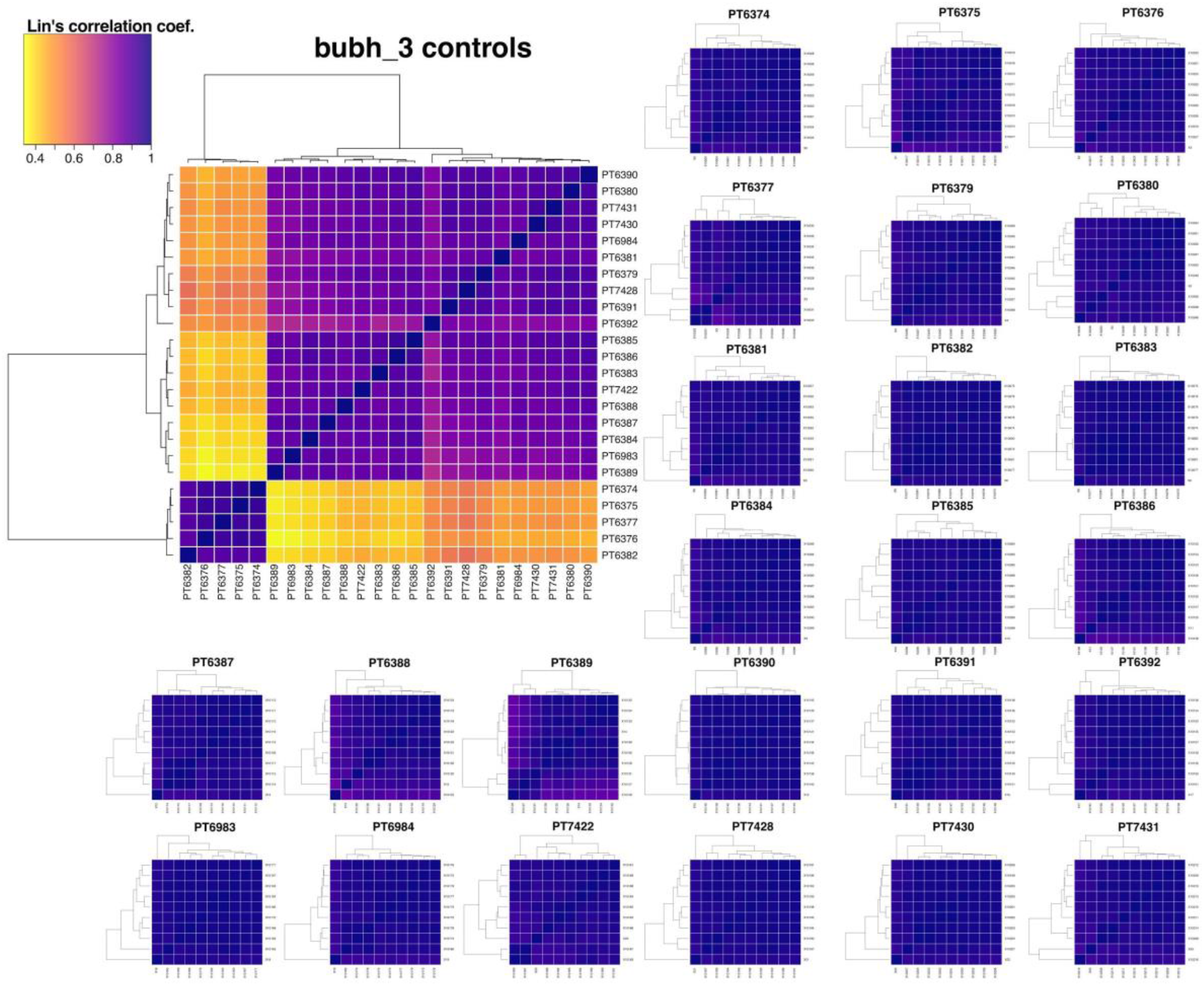
Concordance correlation within 10-plex TMT batches and controls: The heatmaps show hierarchical clustering results for the TMT^10^ −126 (control bubh_3 iPSC line) on the top left, along with the within experiment correlation for all the 24 10-plex TMT batches.

Figure 3 shows that the concordance correlation coefficient within each 10-plex TMT batch is very high, (median value ~ 98% concordance), highlighting the precision of the quantitation within each single batch. However, when the same calculation is applied to the technical replicates of the control iPSC line across the 24 respective batches, the median concordance coefficient drops to ~85%.

To explore this situation further, we calculated the geometric COV (gCOV) for the log10 protein copy numbers^20^ (see methods), both within each 10-plex TMT batch, and across all the 24 controls (Fig. 4a). When we calculated the protein gCOV exclusively within each 10-plex TMT batch, the median was ~1.92%, with all 10-plex TMT batches showing a median protein gCOV < 4%. Accordingly, the data show that for every batch, proteins with a gCOV >10% were considered outliers (Fig. 4a).

**Figure 4.**
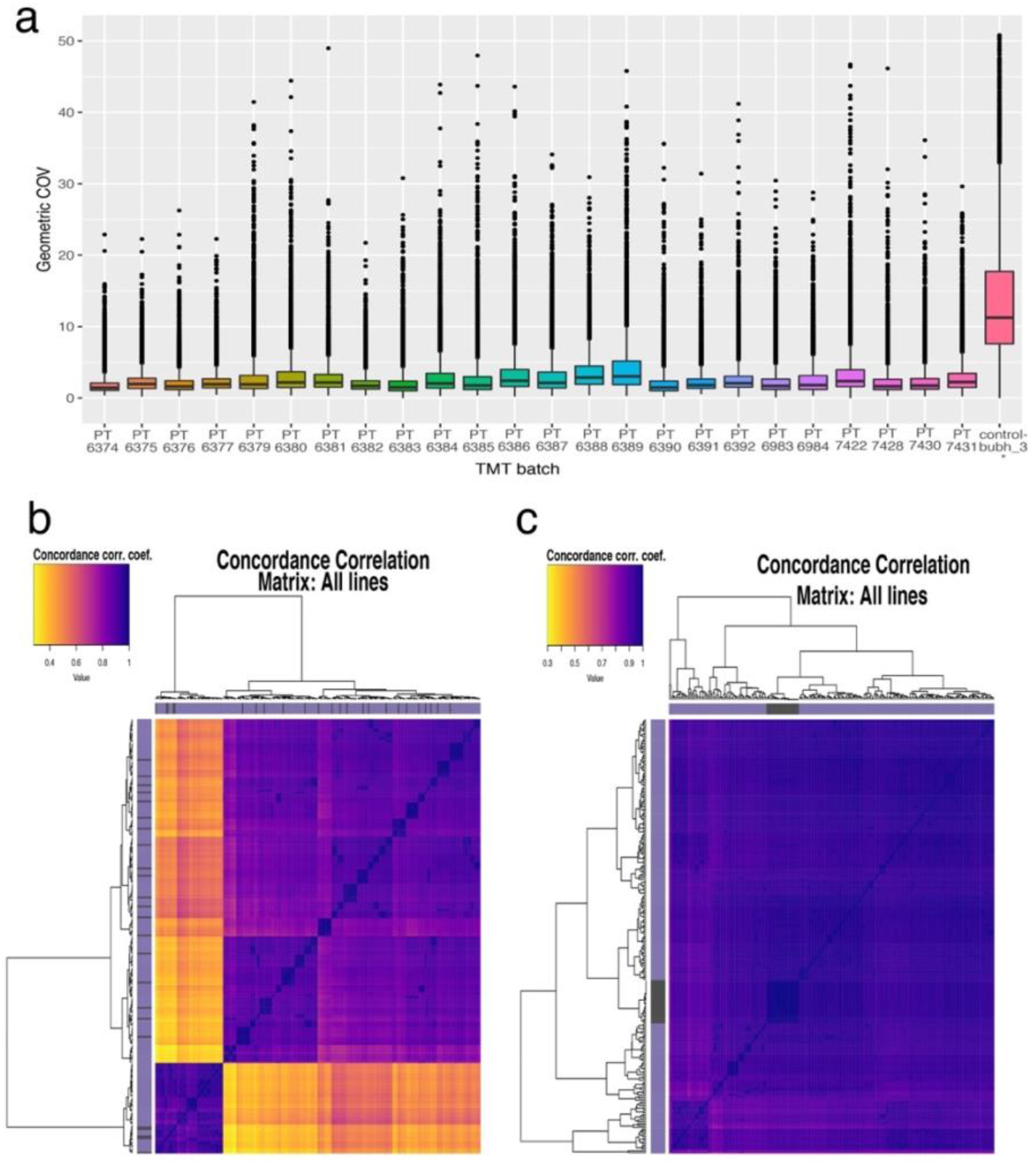
Variation and normalisation: (a) Tukey box plots showing the gCOV for all proteins detected in each 10-plex TMT batch as well as all proteins detected in all the technical replicates (TMT channel 126C in all batches). (b) Hierarchical clustering of all iPSC data using concordance correlation calculated using the log10 raw protein copy numbers (see methods) (c) Hierarchical clustering of all iPSC data using concordance correlation calculated using the log10 control normalised protein copy numbers.

Similarly, we calculated the gCOV for all technical replicates of the control iPS cell line, bubh_3, which were analysed in channel TMT^10^ 126 in every 10-plex TMT batch. The median gCOV of all the proteins detected in the technical replicates, was ~11.28%, i.e. 5.9-fold higher than the median within-batch gCOV. Thus, >50% of all proteins in the technical replicates would be considered outliers in all of the within-batch 10-plex TMT analyses. One example of a protein which highlights this situation is the transcriptional adapter 2-alpha, whose expression levels in the technical replicates of bubh_3 ranged from ~100 copies in PT7428, to ~100,000 copies in PT6374. The gCOV within batch PT6374 is 0.93, and the gCOV across the bubh_3 technical replicates is 24.9.

We note that while the majority of iPSC lines in this study come from healthy donors, some of these TMT batches, e.g. PT6390, contain mixtures of iPSC lines derived from both healthy and donors with rare genetic diseases, including “Usher syndrome”, “Monogenic Diabetes” and “Bardet-Biedl syndrome”. Nonetheless, the median protein gCOV within PT6390 is still ~10 fold lower than the gCOV obtained from analysing the 24 technical replicates.

Our results highlight that while multiplex TMT is a useful and precise methodology for quantitative proteomics, it is important to be aware also of its potential limitations, particularly when analysing data from multiple TMT batches. This is illustrated here by hierarchical clustering of the concordance correlation coefficients for all iPSC data. In this case, 83% of the values for the theoretically identical technical replicates do not cluster together (Fig. 4b). These findings underline that when conducting very large-scale proteomics analyses across multiple separate TMT batches, it is essential to be aware of the potential for batch variation to affect data quality. To reduce the effect of batch variation requires the inclusion of an internal standard in every multi-plex batch to allow for objective data normalisation. For example, here we used 24 technical replicates of a control iPSC line to control for variation between batches (Fig. 4c) (see methods). Using this normalisation method provided a median gCOV of ~2.19% across all cell lines and technical replicates, making the results comparable to the metrics obtained for each individual within-batch analysis. Finding an internal standard that is representative and reproducible is thus vital for multi-batch TMT studies.

### Channel leakage with TMT batches

The iPSC dataset^17,21^ provided us with an excellent opportunity to study the widely recognised effect on data quality and quantitation of both co-isolation interference (CII) and reporter ion interference (RII) within a multi-plex TMT batch. The study utilised iPSC lines derived from both male and female donors within twenty-two of the twenty-four 10-plex TMT batches analysed here. Since only the lines from male donors should include proteins encoded by genes exclusively on the Y chromosome, this provided effectively a set of endogenous “spike-in” peptides, which we could use to monitor both RII between TMT channels, as well as CII.

The dataset detected 13 proteins that were mapped to the Y chromosome. Correspondingly, all peptides derived uniquely from these proteins should only be present in the TMT channels with male cell lines and, in theory, should be absent in the TMT channels with female cell lines. To avoid mismatches arising from shared peptides, we focussed our analysis on a subset of 102 peptides that mapped uniquely to the following Y chromosome specific genes; “DDX3Y”, “EIF1AY”, “KDM5D”, “NLGN4Y”, “RPS4Y1”, “RPS4Y2”, “TBL1Y”, “USP9Y”, “UTY”. Additionally, since two of the 10-plex TMT batches analysed (PT6384 and PT7422) had only female cell lines, any potential Y chromosome-specific peptides that were detected in these batches were treated as potential outliers and discarded from further analysis. Furthermore, batch PT6388 was also considered an outlier and not included for further analysis (see methods). As a result, we focussed on 76 unique, Y chromosome encoded peptides that were used as “male-specific” spike-in references for the analysis of cross-channel RII in the 21 TMT batches that contained mixtures of both male and female derived iPSC lines.

Next, we evaluated how frequently the respective female TMT channels were quantifying signal from Y chromosome-specific peptides (Fig. 5).

**Figure 5.**
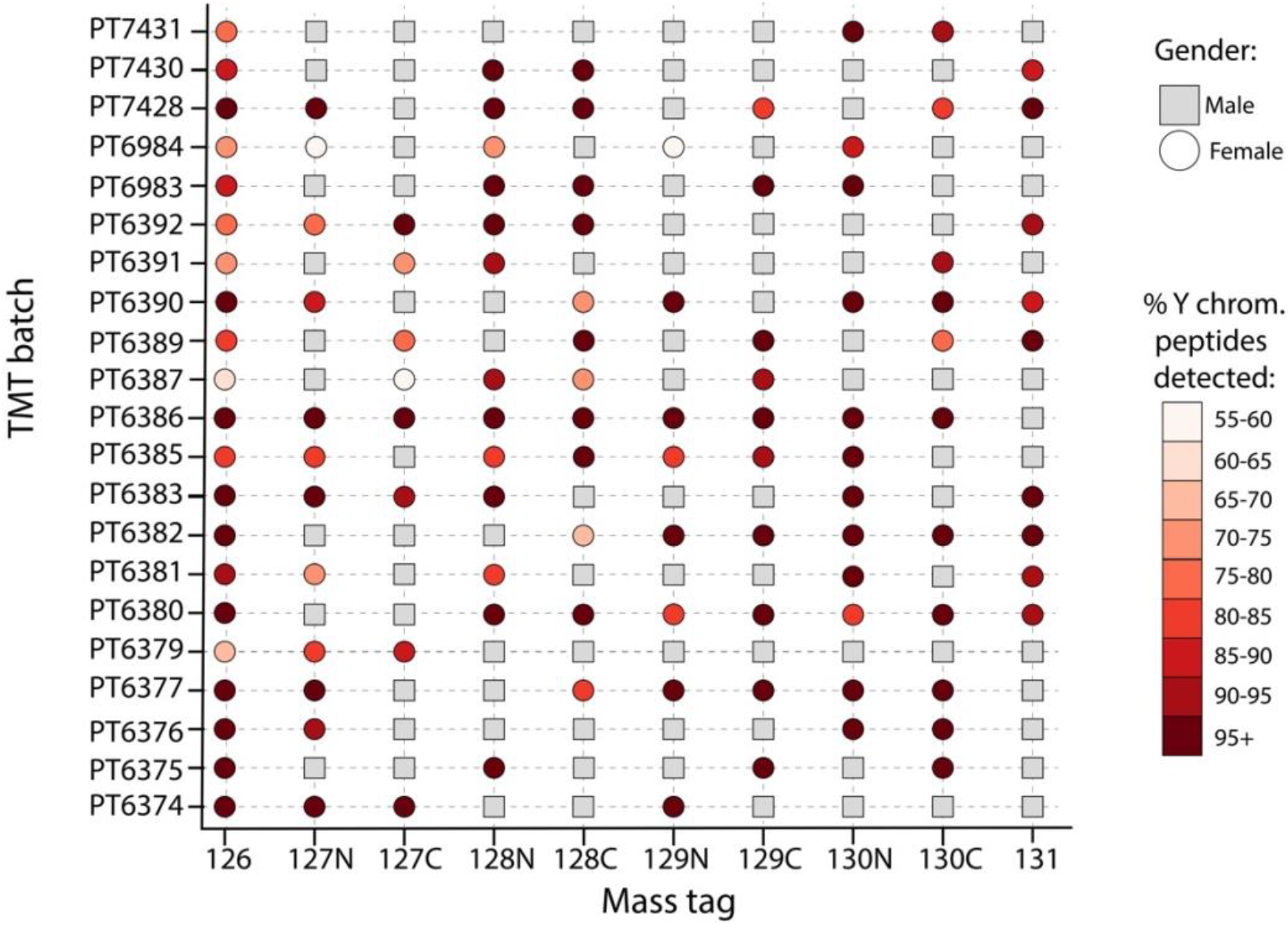
Y chromosome peptides in female channels: Scatter plot showing the gender across all 21 nonoutlier TMT batches and their reporter ion mass tags. Male cell lines are shown as a grey square, female cell lines are represented by a circle. The total number of Y chromosome peptides per batch was calculated and used to determine the percentage of these peptides that were detected within each female channel. The female lines are coloured by this percentage.

Surprisingly, this showed that in all twenty-one 10-plex TMT batches considered here and in all reporter channels containing a female cell line, a minimum of >55% of the Y chromosome-specific peptides identified within the batch also had signal. Remarkably, across all these batches, a median of ~95% of Y chromosome-specific peptides detected in each batch had signal in TMT channels that contained female cell lines. We infer that the appearance of signal for Y chromosome-specific peptides in the channels containing female cell lines likely results from a combination of CII and signal leakage between the TMT channels caused by RII.

We proceeded to evaluate the difference in median-normalised MS3 reporter intensities between male and female lines, across all the twenty-one 10-plex TMT batches (Fig. 6a). Interestingly, the data showed significant variation between batches. For example, some batches, such as PT6380 and PT6386, have ~32-fold difference between male and female peptides, simplifying the detection of false positives due to CII. However, other batches, e.g. PT7431 and PT6391, only show a ~2-fold difference, making the detection of the false positives problematic. We note both of the previously mentioned batches display low Y specific peptide intensities and hence low signal-to-noise ratios, making them more vulnerable to CII.

**Figure 6.**
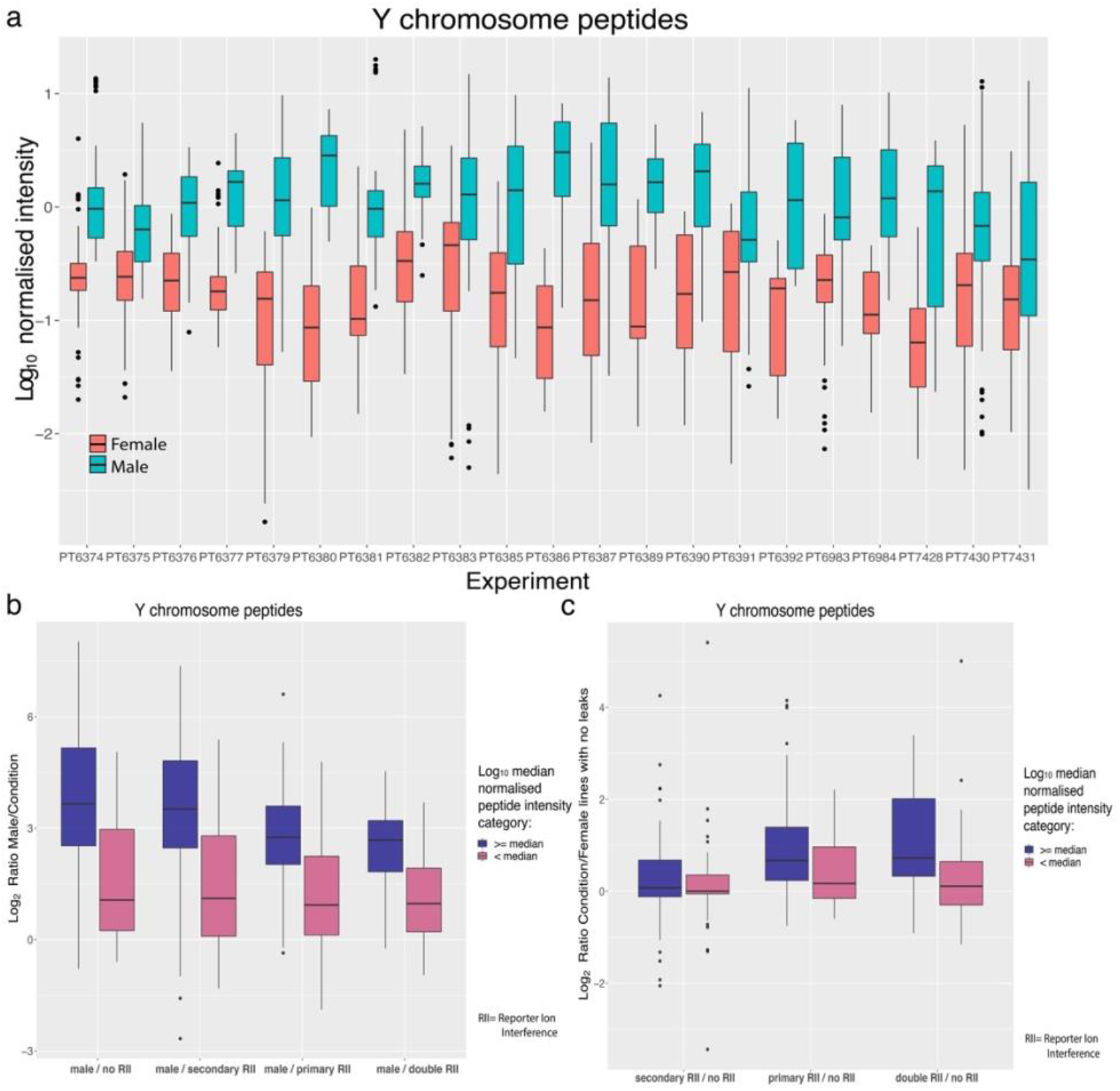
TMT channel leakage analysis: (a) Box plot showing the median normalised intensity of Y chromosome specific peptides detected for both female and male cell lines across all 21 TMT batches. (b) Box plot of ratios for Y chromosome specific peptides, stratified by the median log10 normalised intensity. (c) Box plot of ratios for Y chromosome specific peptides, stratified by the median log10 normalised intensity, comparing leakage conditions vs channels without leaks.

To evaluate co-isolation ion interference (CII), we selected female channels with no primary or secondary RII (see methods), as likely examples of CII^22^. Peptides in male channels show a median of ~8.27-fold higher intensity compared to female channels not affected by RII. However, the effects vary depending on the peptide intensity thresholds. Thus, peptides where the median intensity across male lines was greater than or equal to the global median (−0.20), displayed ~12.5-fold higher intensities. Peptides where the median intensity of male lines was <−0.20 only displayed ~2.09-fold increased intensity, revealing higher vulnerability to CII.

We also examined the potential effects of RII. For this analysis, we calculated a peptide specific ratio for each condition (i.e. “males/double RII”, “males/primary RII”, “males/secondary RII” & “males/no RII”) within each TMT batch and used all of these ratios to generate box plots (Fig 6b). Peptides with a median male intensity <−0.20 were less affected by RII, which is highlighted in Fig. 6b. The male lines were a median of ~2.08 fold higher than female channels not affected by RII, and ~1.95 fold higher than female channels subjected to “primary and secondary RII” (double RII). The female channels subjected to double RII show almost no difference to the female channels not subjected to RII, only ~8% increased intensity. This shows RII had minimal effect on lower intensity peptides.

For peptides with higher intensities (i.e. >=−0.20), we see a reduced effect of CII and increased effect of primary and secondary RII. In this scenario, male lines showed ~12.55-fold higher intensity for Y chromosome-specific peptides than the female channels not subjected to RII and ~6.4-fold higher values than the channels subjected to double RII. In the latter case, we note that the false positives are within the ~8-fold increase/decrease range for *bona fide* changes in protein/peptide expression levels often detected within proteomic datasets^23,24^.

To quantify the differences between primary and secondary RII, we calculated the ratio for each individual peptide within each 10-plex TMT batch showing either primary RII, secondary RII, or double RII (Fig. 6c; see methods). This illustrates that the smallest effect was caused by secondary RII (−1). For high intensity Y chromosome-specific peptides it displayed only a ~1.04-fold increase compared to the channels not affected by RII and virtually no change in low abundance peptides (Fig 6c). The primary RII (+1) produced a more pronounced effect, with a median increase of ~1.58 fold in the high intensity Y chromosome peptides. The combination of primary and secondary RII produced a median ~1.64-fold increase.

These results provide important practical information that aids the design of multi-plex TMT batches to help minimise the potential effects on data quantification of cross condition/population RII.

### Optimising the experimental design

Our data show that the effects of RII should be considered to optimise the experimental design for multi-plex TMT experiments. We advocate for all studies based on more than a single multi-plex TMT batch, a relevant control sample should be included in each batch and assigned to either the 126C, or 127N, channels. These channels avoid primary RII (+1) and are only affected by secondary RII (−1). Secondary RII only causes ~8% increase in intensity, providing better reproducibility for the control. In contrast, placing the control at either the 131N, or 131C channels, increases the impact of RII by exposing the channel to primary RII and thus compromising data quality.

Our results also show TMT experimental designs that can help to minimise the effects of primary and secondary RII between the different populations/conditions. For example, in a 10-plex TMT study, when two conditions are being analysed, each with 5 biological replicates, a 5-5 grouped layout would cause multiple channels to be affected by cross population/condition RII (Fig. 7a). The optimal design would involve alternating the two conditions across the 10 channels (Fig. 7b).

**Figure 7.**
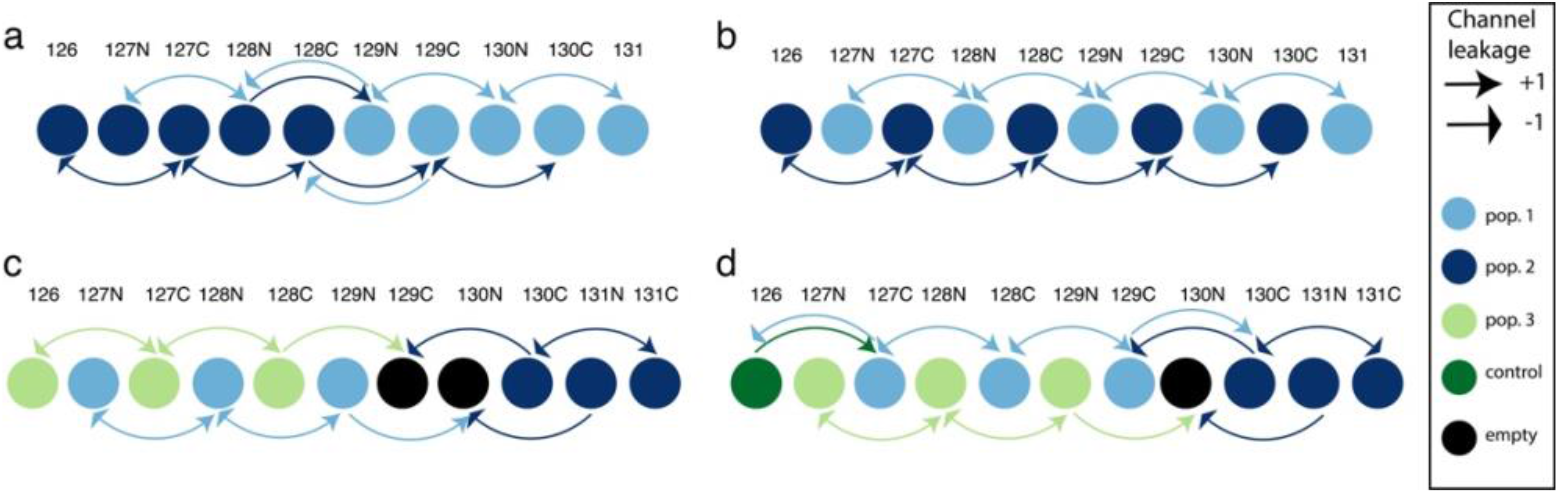
TMT experimental design from reporter ion interference (RII) analysis: (a) 5-5 grouped layout for a 10-plex TMT batch with 5 replicates of two conditions. In this case two channels are being affected by cross population primary and secondary RII. (b) optimal layout for a 10-plex TMT batch with 5 replicates of two conditions with no cross population/condition primary or secondary RII. (c) optimal 11-plex configuration for 3 populations with two empty channels, no channels suffer cross population/condition RII. (d) optimal 11-plex configuration for 3 populations with one empty channel and one control channel. Only two channel suffers primary and secondary RII.

If a control is included, or for studies analysing 3 different conditions in triplicate, (e.g. three time points, or a control and two different perturbations, etc.), we recommend using TMT 11-plex as all 10-plex TMT setups involve increased cross population/condition RII. An 11-plex TMT set up enables a design without RII between the 3 conditions/populations, but requires two empty channels at 129C and 130N to achieve this (Fig. 7c). If a control channel is included, as advocated, then it should be placed in channel 126C, while locating the empty channel to position TMT^11^ 130N, between the alternating experimental conditions and the final replicates of the 3^rd^ condition (Fig. 7d). All of the suggested setups aim to reduce cross condition/population RII as we have shown it decreases quantification accuracy and increases the risk of false positives being detected.

## Discussion

Quantitative proteomic analysis using TMT labelling has become one of the most popular DDA methods currently used, thanks to its multiplexing capabilities, scalability, low missing values index and accuracy when a single multiplexed batch is analysed. However, when very large-scale studies are performed that require the use of multiple parallel TMT batches, the situation becomes more complicated. Here, we have used the analysis of data integrated from 24 separate, 10-plex TMT batches to investigate accuracy, missing values, co-isolation interference, reporter ion interference and experimental design within very large-scale proteomics experiments. We have focussed on a model data set derived from the analysis of human iPS cell lines, derived from both male and female donors^21^.

The resulting data confirm that single batches of multi-plex TMT experiments minimise the typical missing values issue associated in proteomics with Data Dependent Acquisition (DDA), both at the protein and peptide levels. However, this situation changes as data from two or more separate multi-plex TMT batches are integrated. When multiple batches are combined the missing values index inflates rapidly. This effect is particularly striking at the peptide level, where integrating data from only two different batches causes the missing values index to increase from <2% to ~25% Even though the inflation rate at the protein level is lower, the integration of the second batch pushes missing protein values from <0.5% to >10%. This inflationary effect can decrease the accuracy of results derived from large-scale experiments that compare data generated from multiple TMT batches. One potential solution would be to utilise MS2-based TMT quantitation, as this has been reported to produce more total peptide identifications^25^, however there is no guarantee this will reduce peptide/protein missing values across batches and it will intensify the effect of the co-isolation interference.

While single TMT batches can provide remarkably precise results, we found that this result is often not reproducible across batches. To study reproducibility, for every protein we calculated the geometric Coefficient of Variation (gCOV) for the technical replicates of the control line, analysed across 24 separate TMT batches, and the gCOV within each of the twenty-four 10-plex TMT batches. The gCOV of the technical replicates (all derived and reprogrammed from the same donor of course) was ~5.9-fold higher than the median gCOV for iPSC lines derived from different donors analysed within the same 10-plex TMT batch. A recent study of population wide variation within iPSC^26^ revealed the main component driving variation was the donor effect. However, our data suggests unnormalized batch effects can be a bigger issue. This underlines the importance of including a common control sample within each TMT batch to allow for objective data normalisation to minimise the batch effects. We showed that by introducing a suitable control within each TMT batch the data can be normalised effectively, thereby reducing the gCOV to a level that is comparable to the variation seen within a single TMT batch. The challenge lies in identifying a suitable control that is truly representative for most proteins being compared within the experiment and in creating a control that is highly reproducible across all the TMT batches.

This study has also highlighted the issue of reporter ion interference (RII) and co-isolation interference (CII), which are parameters than can compromise data accuracy in TMT experiments. The dataset we selected provided an ideal set up to analyse these factors, as it contained iPSC lines derived from both male and female donors. Thus, by identifying a set of peptides uniquely mapped to the male-specific Y chromosome, these provided a convenient set of internal controls to monitor the expression of false positives. Furthermore, we compared 21 different 10-plex TMT batches with different numbers of male and female derived cell lines, assigned in different channel combinations. We were thus able to determine how the arrangement of channels can be optimised to minimise the impact of RII for different experimental design scenarios^22^.

The data showed that even for 10-plex TMT batches with only two male channels (PT6380), the remaining 8 female channels still had signal for ~95.8% of all the Y chromosome-specific peptides that were detected in that batch. Moreover, low abundance peptides (i.e. normalised intensity lower than the median) were only ~2.09-fold higher intensity in the males than the females. This highlights the potential effects of CII within TMT experiments, particularly for quantitating low abundance peptides. We note that the CII issue has been reduced, though not eliminated, with newer generation Orbitrap MS instruments, where the improved source and quadrupole have enhanced the signal to noise ratio^22^. Furthermore new isobaric tagging methods have been developed which claim to be CII free^27^, however their multiplexing capability is currently limited to a 6-plex.

The data analysed here also highlighted the effects of primary and secondary RII in channels with high intensity peptides. Thus, reporter channels affected by both primary and secondary RII showed a median signal increase ~1.64-fold higher than channels not subjected to RII. To best avoid this situation, we have used these data to propose optimised experimental set ups for assigning samples to specific channels that can either minimize, or eliminate (when possible), the effect of primary and secondary RII between conditions/populations. Nonetheless, we highlight that mixing significantly different populations within a TMT batch, for example iPSCs and terminally differentiated somatic cells, still poses the risk of false positives being identified, as illustrated here by the Y chromosome-specific peptides detected within all female cell lines.

In conclusion, TMT is a valuable methodology for DDA analysis and its potential to increase scalability and precise quantitation have made it a justifiably popular approach for high-throughput proteomic studies. Here, we have provided an in-depth, practical evaluation of parameters affecting the generation of high-quality quantitative data from very large-scale TMT-based proteomics analyses, and we highlight some of the limitations which should be considered when planning these experiments. We hope the resulting information will prove useful for improving experimental design and resulting data quality for many future proteomics projects.

## Methods

### TMT sample processing and LC-MS

The data analysed within this study used a SPS-MS3 method for LC-MS on an Orbitrap Fusion Tribid mass spectrometer (Thermo Fisher). For details regarding sample preparation and LC-MS methods see ^17^.

### Identification & Quantification

The data from all 24 10-plex TMT batches were collected and analysed simultaneously, using Maxquant^28^ v. 1.6.0.13. The FDR threshold was set to 5% for each of the respective Peptide Spectrum Match (PSM) and Protein levels. Proteins and peptides were identified using UniProt (SwissProt & TrEMBL). Run parameters are accessible at ProteomeXchange^29^ via the PRIDE repository ^30^, along with the full MaxQuant^28^ quantification output (PDX010557).

### Copy number generation

Protein copy numbers were calculated following the proteomic ruler approach^31^ and using the MS3 intensity for each protein group identified. These uncorrected copy numbers were used to study the geometric coefficient of variation (gCOV), which will be referred to here as “raw copy numbers”.

To control for technical variation between the 24 different 10-plex batches, a correction factor was applied to adjust the protein copy numbers. This correction factor was derived from analysing the protein levels in the reference iPSC line, which was present in channel 126C on every batch. Specifically, for every protein, a median copy number was calculated using values from all 24 technical replicates. Thus, the protein copy number derived from the control cell line within each TMT batch was divided by the median copy number for that protein, based upon values from all 24 controls. This ratio was then used as a correction factor to normalise the expression values for all proteins detected within each respective 10-plex TMT batch. The corrected values are referred to here as “normalised copy numbers”.

### Missing value calculations

First, to estimate missing values within this DDA analysis, a list of unique proteins/peptides that were detected with at least 1 reporter intensity greater than zero were calculated for each batch. To determine the number of missing values within each 10-plex TMT batch, the number of unique proteins per reporter channel was compared to the number of unique proteins/peptides identified within the batch. This approach was applied to generate the missing value calculations for each of the 24 individual 10-plex TMT batches. To assess the effect of integrating multiple TMT batches, random sampling was performed to estimate how missing values are affected by a progressive increase in the number of 10-plex TMT batches analysed. This was performed in an incremental fashion starting from 2 and finishing with 23 batches, with 500 iterations per level.

At each level a new a list of proteins/peptides detected with at least 1 reporter ion intensity greater than zero within any of the integrated TMT batches was calculated, and the number of proteins/peptides with intensity greater than 0 per reporter channel was evaluated against the new list.

### Coefficient of variation

The geometric coefficient of variation (gCOV) in protein abundance levels was calculated using log_10_ transformed protein copy numbers. These data showed a log normal distribution, therefore the COV was calculated using the geometric method^20^ via the R package “PKNCA”^32^ version 0.8.5.

The protein gCOV within each 10-plex TMT batch was calculated for all 10 cell lines within the same batch, using all proteins detected in every reporter channel. The control gCOV was calculated using proteins that were detected in the TMT^10^ −126C (control) channel across all of the 24 10-plex TMT batches.

### Correlation Clustering

For each 10-plex TMT batch, a concordance correlation value was calculated for all cell lines within the same batch. The calculations were performed using “correlation()” function from the R package “agricolae” version 1.2.8. The heatmaps were generated using “heatmap.2” from the R package “gplots” version 3.0.1 and using hierarchical clustering with “ward.D2” to calculate distances.

The same process was applied to calculate the concordance correlation values for the controls, i.e. using reporter channel 126C in all TMT batches.

### Peptide intensity normalisation

Peptide intensities were median normalised. For this, after filtering out peptides with zero MS3 intensity, a median intensity value was calculated for all reporter channels in each of the 24 10-plex TMT batches. Every reporter intensity greater than zero was divided by the median value for the reporter ion intensity of its specific channel, this data was then log_10_ transformed. For the intensity histogram, a median across all reporter channels and batches was used.

### Reporter ion interference classification

The reporter ion interference (RII) targets are based on a typical product data sheet for 10-plex TMT Label Reagents from ThermoFisher Scientific, as summarised in the table below:

| Mass Tag | Reporter channel | -1 (secondary RII) | +1 (primary RII) |
| --- | --- | --- | --- |
| TMT<sup>10</sup> -126 | 0 | -- | 127C |
| TMT<sup>10</sup> -127N | 1 | -- | 128N |
| TMT<sup>10</sup> -127C | 2 | 126 | 128C |
| TMT<sup>10</sup> -128N | 3 | 127N | 129N |
| TMT<sup>10</sup> -128C | 4 | 127C | 129C |
| TMT<sup>10</sup> -129N | 5 | 128N | 130N |
| TMT<sup>10</sup> -129C | 6 | 128C | 130C |
| TMT<sup>10</sup> -130N | 7 | 129N | 131 |
| TMT<sup>10</sup> -130C | 8 | 129C | -- |
| TMT<sup>10</sup> -131 | 9 | 130N | -- |

### Analysis of Channel Leakage

To study the effect of leakage across different channels, we selected a subset of 102 peptides that were specific to the following list of protein coding genes uniquely located on the Y chromosome; “DDX3Y”, “EIF1AY”, “KDM5D”, “NLGN4Y”, “RPS4Y1”, “RPS4Y2”, “TBL1Y”, “USP9Y” & “UTY”.

This approach of using peptide values from Y chromosome specific genes depends upon there being a diverse mixture of male and female donor-derived iPSC lines in each 10-plex TMT batch. However, two of the 24 TMT batches comprised exclusively female donor-derived iPSCs, which had been shown not to have Y chromosome derived DNA in QC analyses^21^. For these female donor-specific batches, any peptide assigned to Y chromosome specific genes was excluded from the analysis. An additional batch, PT6388, identified only 1 Y chromosome-specific peptide which displayed an irregular behaviour, and was hence also discarded from the analysis. A final subset of 76 Y chromosome-specific peptides were used for this analysis (see supplemental data for list).

### Peptide ratios

The peptide ratios across multiple reporter ion interference conditions (“primary” or “+1 leak”, “secondary” or “−1 leak”, “primary and secondary” or “double leak” and “no leak”) were calculated within each 10-plex TMT batch, utilising the median-normalised log10 MS3 reporter intensities. The ratios were calculated for each individual peptide within each 10-plex TMT batch. The box plot showing peptide ratios utilised all these calculated ratios, plotted using “ggplot2” version 3.0.0^33^.

## Author contributions

A.I.L. supervised the project. J.H. assisted in the interpretation and analysis of the data. D.B. assisted in the interpretation of the data. A.B. conceived the study and performed the data analysis and interpretation. The paper was written by A.B. and edited by all authors.

## Data availability

The raw files used for this analysis are accessible at ProteomeXchange^29^ via the PRIDE repository ^30^, along with the full MaxQuant^28^ processed output (PDX010557).

## Acknowledgments

This works was funded by the Wellcome Trust /MRC [098503/E/12/Z] and Wellcome Trust grants [073980/Z/03/Z, 105024/Z/14/Z].

